# An autoregulatory switch in sex-specific *phf7* transcription causes loss of sexual identity and tumors in the *Drosophila* female germline

**DOI:** 10.1101/2020.05.13.093344

**Authors:** Anne E. Smolko, Laura Shapiro-Kulnane, Helen K. Salz

## Abstract

Maintenance of germ cell sexual identity is essential for reproduction. Entry into the spermatogenesis or oogenesis pathway requires that the appropriate gene network is activated and the antagonist network is silenced. For example, in *Drosophila* female germ cells, forced expression of the testis-specific PHD finger protein 7 (PHF7) disrupts oogenesis leading to either an agametic or germ cell tumor phenotype. Here we show that PHF7 expressing ovarian germ cells inappropriately express hundreds of genes, many of which are male germline genes. We find that the majority of genes under PHF7 control in female germ cells are not under PHF7 control in male germ cells, suggesting that PHF7 is acting in a tissue-specific manner. Remarkably, transcriptional reprogramming includes a positive autoregulatory feedback mechanism in which ectopic PHF7 overcomes its own transcriptional repression through promoter switching. Furthermore, we find that tumorigenic capacity is dependent on the dosage of *phf7*. This study reveals that high levels of ectopic PHF7 in female germ cells leads to a loss of sexual identity and promotion of a regulatory circuit beneficial for tumor initiation and progression.

## INTRODUCTION

Germ cell development culminates in the production of sexually dimorphic haploid gametes: sperm and eggs. In most animals, cells destined to become germ cells are set aside during embryogenesis and migrate to the developing gonad. There they exhibit sex specific division rates and gene expression programs, ultimately leading to meiosis and differentiation into morphologically and functionally distinct gametes (Lesch and Page, 2012). Germ cell development is not possible when the sexual identity of the germ cells and the surrounding somatic gonadal cells do not match (Salz et al., 2017). Successful reproduction, therefore, requires that the appropriate sex-specific expression network be activated and the antagonist network be silenced.

In *Drosophila melanogaster* germ cells, the female/male decision is initially guided by the sex of the developing somatic gonad (Casper and van Doren, 2009; Hashiyama et al., 2011; Horabin et al., 1995; Staab et al., 1996; Wawersik et al., 2005). Extrinsic control is eventually lost, and sexual identity is maintained by cell-intrinsic mechanisms (Casper and van Doren, 2009). In female germ cells, maintenance of the embryonic sex fate decision requires the female-specific RNA binding protein Sex-lethal (SXL) (Chau et al., 2009; Schüpbach, 1985; Shapiro-Kulnane et al., 2015; Smolko et al., 2018). When germ cells lack SXL protein, differentiation is blocked and germ cell tumors are formed. Although loss of SXL leads to the ectopic sex-inappropriate transcription of hundreds of genes, dysregulation of one spermatogenesis gene, *PHD Finger Protein 7 (phf7)*, was found to be a major driver of the germ cell tumor phenotype (Shapiro-Kulnane et al., 2015).

PHF7 is a predicted chromatin reader that was first identified in a screen for genes expressed in male but not female embryonic germ cells (Yang et al., 2012). In the adult testes, protein expression is restricted to the nuclei of the undifferentiated germline stem cells and early spermatogonia (Yang et al., 2012; Yang et al., 2017). However, loss of *phf7* has only minimal impact on spermatogenesis. Mutant males harbor fewer differentiating spermatogonial cysts, resulting in fewer progeny than wild-type. Reduced fecundity appears to be caused by a failure to control expression of a small set of male germ cell genes (Yang et al., 2017).

Although not essential for germ cell development in males, it is crucial to prevent PHF7 expression in female germ cells. PHF7-expressing ovarian germ cells fail to differentiate, resulting in an agametic or germ cell tumor phenotype (Shapiro-Kulnane et al., 2015; Yang et al., 2012). Interestingly, when PHF7-expressing XX germ cells develop in a sexually transformed somatic environment, they can produce sperm, albeit at a low frequency (Yang et al., 2012). These studies suggest that ectopic PHF7 expression is able to drive the germ cell towards a male developmental program. However, the impact of ectopic PHF7 on the transcriptional landscape is not known.

In this work, we combined genetic and genomic approaches to understand the consequences of ectopic PHF7 expression in ovarian germ cells. As expected, we find that a female to male identity switch underlies tumor formation. However, the majority of genes under PHF7 control in ovarian germ cells are not under PHF7 control in male germ cells. This suggests that PHF7 affects gene expression in a tissue-specific manner. Ectopic transcriptional reprogramming activity includes a positive autoregulatory feedback mechanism in which PHF7 can overcome transcriptional repression of other *phf7* copies in the genome. The resulting increase in PHF7 expression correlates with an increase in tumorigenic capacity. Lastly, we show that transcriptional autoregulation and oncogenic properties of PHF7 requires the PHD fingers. Together, our work supports a model in which PHF7 reprograms transcription in the female germline by redirecting chromatin remodeling complexes to inappropriately activate male germ cell genes; underscoring the importance of preventing expression of lineage-inappropriate genes for maintaining tissue homeostasis.

## RESULTS AND DISCUSSION

### Deletion of the PHD fingers creates an inactive *phf7* allele

*phf7* encodes a 520 amino acid protein with three adjacent N-terminal PHD fingers: a canonical PHD zinc finger domain (ZNF_PHD), an extended PHD (ePHD) domain, and a RING-finger domain (ZNF_RING) (Mitchell et al., 2019). PHD finger proteins are often involved in chromatin and transcriptional regulation. To test whether deleting this region impacts function *in vivo*, we used the tissue-specific GAL4/UAS induction system to force expression of *phf7* cDNA carrying an in-frame deletion of amino acids 68-111 (UASz::*phf7*^*ΔPHD*^). This transgene and a control wild-type transgene (UASz::*phf7)* were inserted into the genome via site-specific recombination in to the same location to allow direct comparisons (Fig. 1A).

**Fig. 1.**
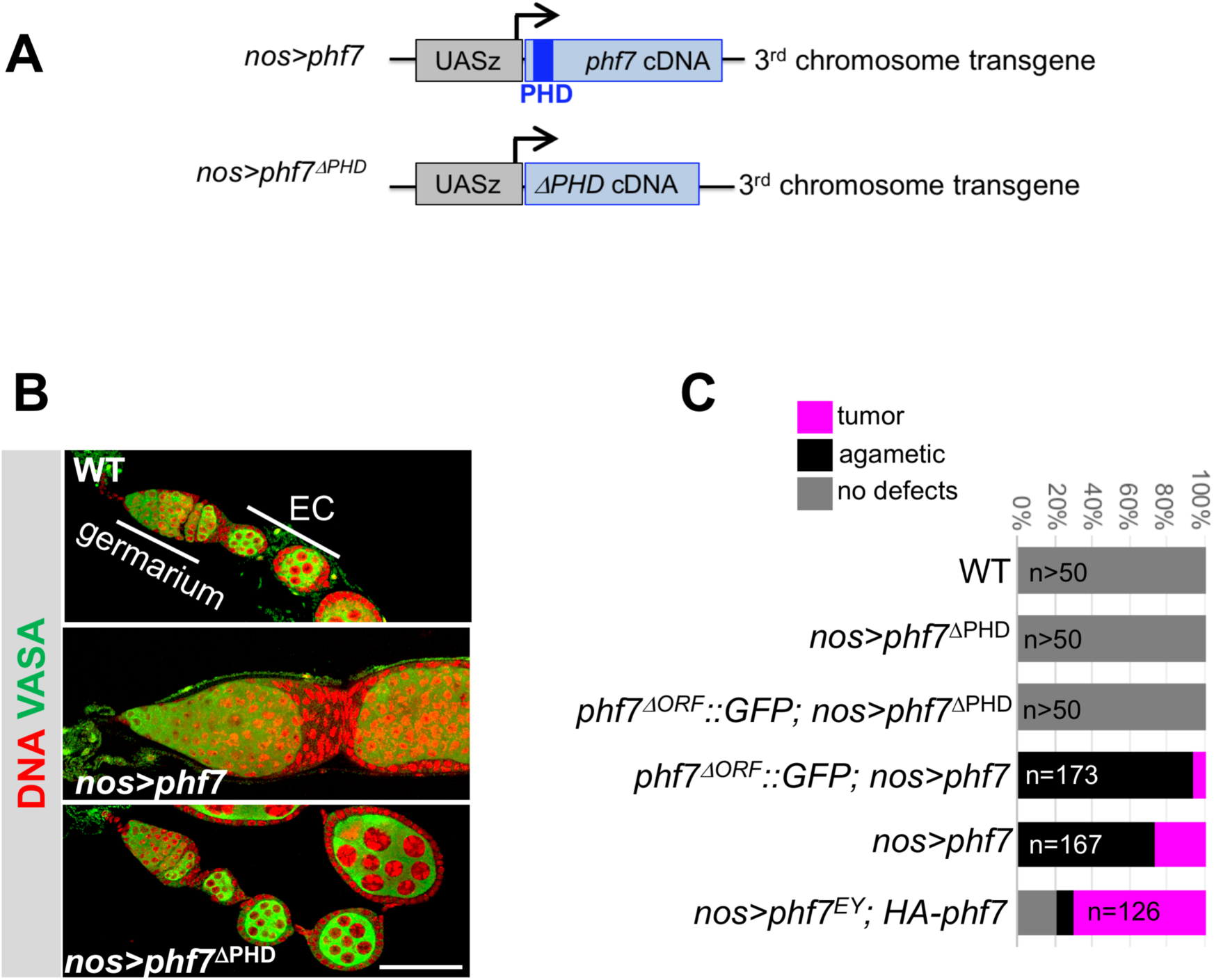
Deletion of the PHD domain disables ectopic PHF7 activity in female germ cells. (A) Schematic representation of the fly lines used to express wild-type and mutant *phf7* cDNAs via the GAL4/UAS system. The cDNAs were cloned into the UASz expression vector and inserted into the third chromosome 65B2 attP site. (B) Ectopic expression of wild type (*nos*>*phf7)* but not mutant *(nos*> *phf7*^*ΔPHD*^) PHF7 in germ cells disrupts oogenesis. Representative confocal images of mutant and control ovarioles, including the germarium and egg chambers (EC), stained for Vasa (green) and DNA (red). Scale bar, 50 μm. (C) Quantification of the different mutant ovariole phenotypes observed in this study. The number of ovarioles scored is indicated.

Using the germ cell specific driver, *nos-Gal4::VP16*, we found that expression delivered from the wild-type UASz-*phf7* transgene caused females to be sterile. Progression of egg chamber formation was analyzed by staining for DNA and Vasa, a germ cell specific marker (Fig. 1B). The fly ovary is composed of 16-20 ovarioles, each of which is organized as an assembly line that generates egg chambers (Hinnant et al., 2020). At the anterior end of the ovariole, a structure called the germarium houses two to three germline stem cells. Each stem cell divides to create one daughter cell that remains a stem cell and a second daughter cell that enters the differentiation pathway. Differentiation begins with 4 incomplete mitotic divisions to generate a 16-cell cyst that develops into an egg chamber containing 1 oocyte and 15 supporting nurse cells. As expected, the ovaries of *nos*> UASz-*phf7* mutant females exhibited morphological defects. 74% (n=167) of *nos*> UASz-*phf7* mutant ovarioles were agametic. In the remaining 26% of the mutant ovarioles we observed a tumor phenotype (Fig. 1B, C) defined by the accumulation of excess germ cells in the germarium and the failure to form egg chambers with an oocyte and nurse cells.

In sharp contrast, the *nos*> UASz-*phf7*^*ΔPHD*^ females were fertile and their ovaries were similar to wild-type (Fig. 1B, C). The failure to generate a mutant phenotype in this ectopic expression assay indicates that deleting the PHD finger domain inactivates *phf7.* Overall, these data show that ectopic expression in female germ cells is sufficient to disrupt oogenesis and suggests that the PHD finger domain is required. However, we cannot rule out other explanations for the loss of ectopic *phf7*^*ΔPHD*^ activity, such as protein instability, because antibodies against PHF7 are unavailable.

### Ectopic Phf7 functions in a feedback loop

Autoregulatory feedback mechanisms are often used to maintain expression of fate determining genes (Crews and Pearson, 2009). Therefore, we asked whether forced Phf7 expression from the transgene could induce testis-like expression at the endogenous locus.

To monitor expression at the endogenous locus we used CRISPR to replace the open reading frame with GFP (*phf7*^*ΔORF*^::*GFP;* Fig. 2A). We determined that the reporter does not interfere with female fertility and that it recapitulates PHF7’s sex-specific RNA and protein expression patterns (Fig. S1). When ectopic *phf7* expression (nos> UASz-*phf7*) is induced in this background, GFP protein is detected in tumors (Fig. 2B). We conclude that ectopic PHF7 stimulates testis-like protein expression from an edited allele at the endogenous locus. In contrast, we did not observe GFP staining when ectopic expression of the inactive *phf7*^*ΔPHD*^ allele was induced in this background *(phf7*^*ΔORF*^::*GFP; nos*> *UASz-phf7*^*ΔPHD*^; Fig. 2B), demonstrating that functional PHF7 protein is required for transactivation.

**Fig. 2.**
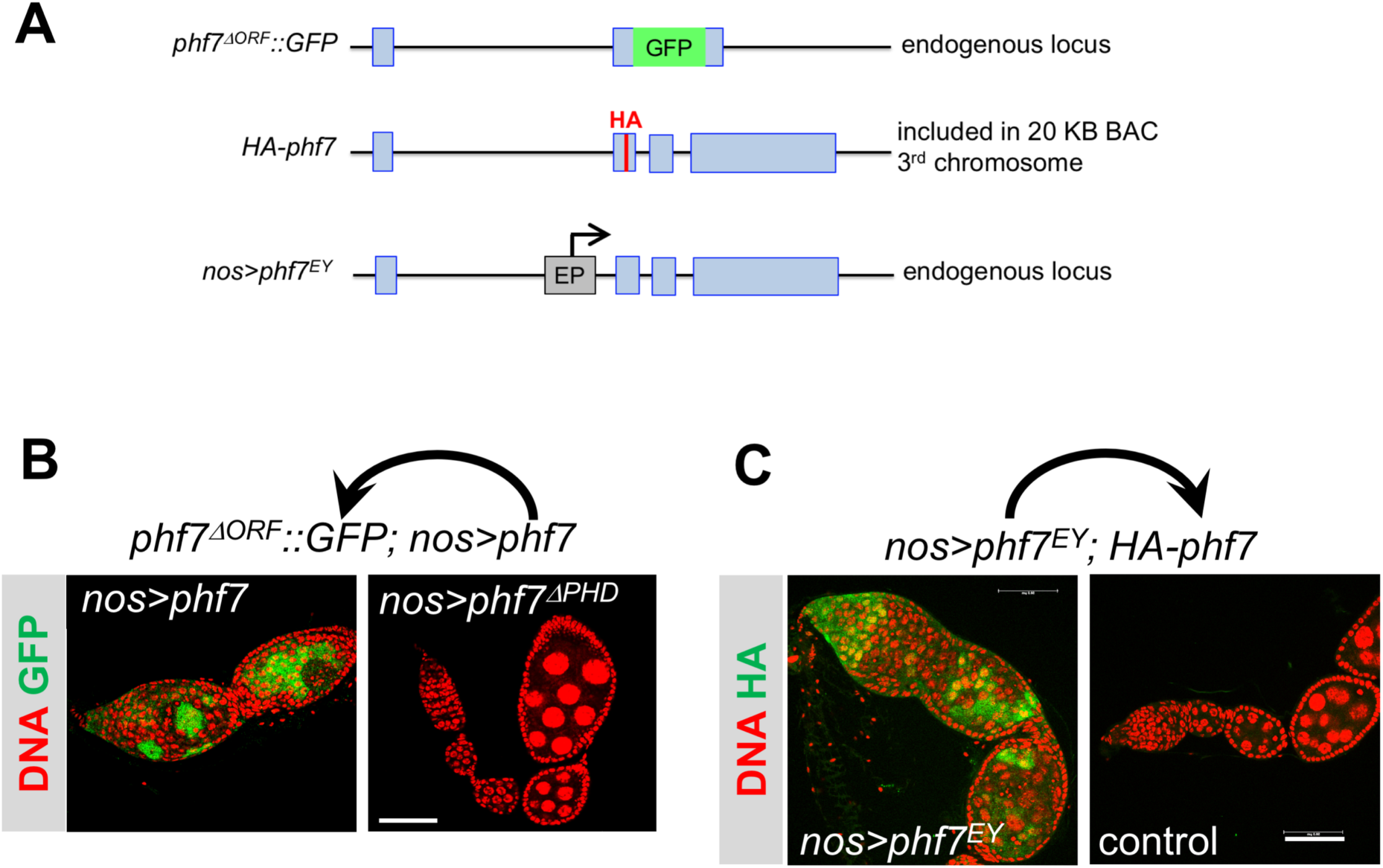
Ectopic PHF7 stimulates testis-like expression from reporter alleles. (A) Schematic representations of Top: the knock-in *phf7*^*ΔORF*^::*GFP* allele used to report on *phf7* activity. Middle: the 3^rd^ chromosome HA-tagged *phf7* reporter allele, located within a 20 kb BAC inserted into the 65B2 attP site. Bottom: the *phf7*^*EY*^ allele in which the UASp containing P{EPgy2} element inserted into the first intron of the endogenous locus is used to ectopically express *phf7* via the GAL4/UAS system. (B) Ectopic PHF7 can induce testis-like expression from the knock-in *phf7*^*ΔORF*^::*GFP* allele. Confocal images of ovarioles from *phf7*^*ΔORF*^::*GFP*; *nos*>*phf7* and control *phf7*^*ΔORF*^::*GFP*;*nos*>*phf7*^*ΔPHD*^ females stained for GFP protein (green) and DNA (red). Scale bar, 50 μm. (C) Ectopic PHF7 can induce testis-like expression from a 3^rd^ chromosome HA-tagged reporter construct. Confocal images of ovarioles from mutant *nos*>*phf7*^*EY*^; *HA-phf7* and control *nos; HA-phf7* females stained for HA (green) and DNA (red). Scale bar, 50 μm.

Next, we sought to demonstrate positive autoregulation using a different genetic paradigm. We forced PHF7 expression from the endogenous locus via an UASp-containing EP transposable element insertion, *P{EPgy2}phf7*^*EY03023*^ (*phf7*^*EY*^) located within the first intron (Fig. 2A). Driving expression of *phf7*^*EY*^ with the germline *nos-GAL4* driver has been shown to drive tumor formation, albeit at a low frequency (Shapiro-Kulnane et al., 2015; Yang et al., 2012). We assessed transactivation with a HA-tagged *phf7* locus embedded in a 20 kb BAC rescue construct located on the 3^rd^ chromosome (Fig. 2A). This transgene has been shown to serve as a faithful reporter of PHF7’s sex-specific protein expression pattern (Shapiro-Kulnane et al., 2015; Yang et al., 2012). When these two genetic elements are combined (*nos*>*phf7*^*EY*^; *HA-phf7*), ectopic HA-PHF7 protein is detected in tumors (Fig. 2C). We conclude that ectopic PHF7 stimulates testis-like protein expression from the transgenic tagged copy of *phf7.* This shows that ectopic PHF7 can stimulate testis-like expression from any *phf7* allele.

Together this data shows that once expressed in ovarian germ cells, PHF7 can increase its own expression via a positive autoregulatory feedback loop. This finding suggests that there may be a correlation between copy number and phenotype. In this context it is interesting to note that in the absence of functional *phf7* copies at the endogenous locus (*phf7*^*ΔORF*^::*GFP; nos*>*UASz-phf7*), the frequency of tumor formation upon induction with a third chromosome transgene was only 6%, and the majority of the mutant ovarioles were agametic (Fig. 1C). Two functional copies at the endogenous locus, using the same third chromosome transgene to ectopically express *phf7*, shifted the distribution of the mutant phenotypes towards a germ cell tumor phenotype (26% tumors in *nos*>*UASz-phf7*). Finally, using a different genetic paradigm with three full length copies of *phf7* dramatically increased the penetrance of the tumor phenotype to 70% (*nos*>*phf7*^*EY*^; *HA-phf7*). We therefore conclude that the level of PHF7 protein dictates the phenotypic outcome, with the highest levels required for tumor initiation and progression.

### Ectopic PHF7 transactivates via promoter-switching

Although PHF7 protein is normally limited to male germ cells, *phf7* mRNAs are expressed in both male and female germ cells (Fig. 3A). Sex-specific regulation is achieved by a mechanism that relies primarily on alternative promoter choice and transcription start site (TSS) selection. In ovaries, transcription from the downstream TSS produces an mRNA, *phf7-RA*, but no protein is detectable. In testis, transcription from the upstream TSS produces a longer translatable mRNA, called *phf7-RC* (Shapiro-Kulnane et al., 2015). Our discovery that ectopic PHF7 can stimulate protein expression from any *phf7* allele suggests a mechanism that includes transcriptional switching to the male-specific TSS. With the identification of genetic conditions that increased the penetrance of the tumor phenotype to 70% (*nos*>*phf7*^*EY*^; *HA-phf7*), we were able to test this hypothesis by assaying for the presence of the testis-specific *phf7-RC* RNA isoform.

**Fig. 3.**
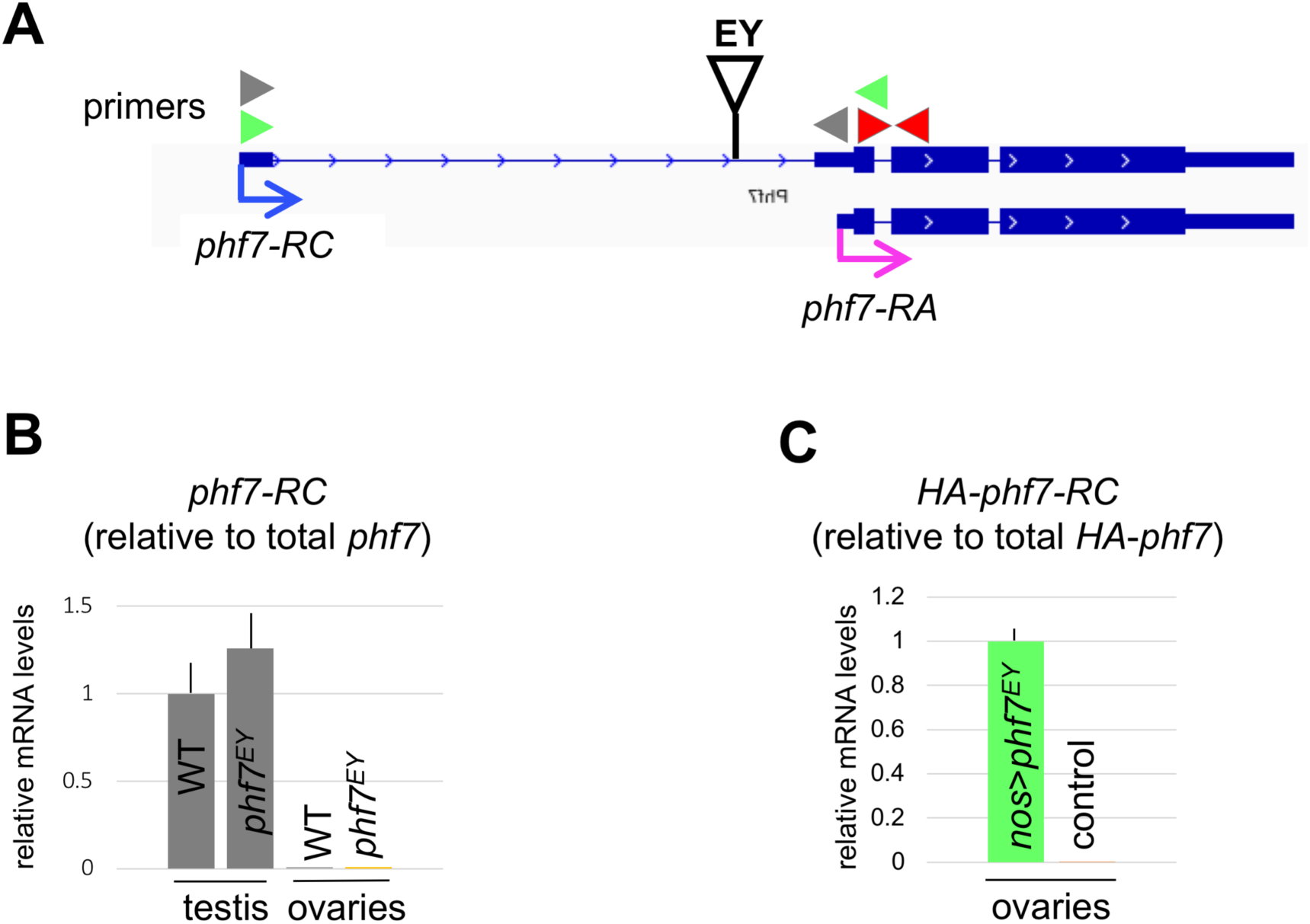
Ectopic PHF7 transactivates via promoter-switching. (A) IGV genome browser view of the two major *phf7* transcripts, the testis-specific *phf7-RC* (blue arrow) and *phf7-RA* (pink arrow). Exons are represented by blue blocks connected by horizontal lines representing introns. The insertion site of the UAS-containing insertion in *phf7*^*EY*^ is represented by a triangle. Location of primers for RT-qPCR are indicated by arrowheads. (B) RT-qPCR measurements of *phf7-RC* transcript in ovaries and testis from wild-type and *phf7*^*EY*^ mutant animals (grey primer pairs). Expression is normalized to the total level of *phf7* (red primer pairs). Error Bars indicate standard deviation of three biological replicates. (C) RT-qPCR measurements of transgenic *HA-phf7-RC* transcript in ovaries from mutant *nos*>*phf7*^*EY*^; *HA-phf7* and control *nos; HA-phf7* females. Primers to the HA tag, located at the beginning of the open reading frame were used to distinguish transgenic *phf7* RNA from the endogenous products: green primer pairs for *HA-phf7-RC* and modified red primer pairs for total *HA-phf7.* Error Bars indicate standard deviation of three biological replicates.

Using RT-PCR, we found that in control *phf7*^*EY*^ ovaries, the EP insertion by itself does not interfere with *phf7’s* sexually dimorphic transcription pattern (Fig. 3B). Furthermore, we found that transcription from the transgenic HA-tagged *phf7* locus is also regulated appropriately as no HA-tagged *phf7-RC (HA-phf7-RC)* transcript is detected in ovaries (Fig. 3C, control). In *nos*>*phf7*^*EY*^; *HA-phf7* mutant ovaries, however, we found that *HA-phf7-RC* is ectopically expressed (Fig. 3C). This work demonstrates that ectopic PHF7 stimulates testis-like transcription from the transgenic, tagged copy of *phf7.*

In agreement with our RT-PCR analysis, alignment of RNA-sequencing (RNA-seq) data showed that the testis-specific *phf7-RC* transcript is ectopically expressed in *nos*>*phf7*^*EY*^; *HA-phf7* mutant ovaries (Fig. 4A). These data also illustrate that novel RNAs are produced from the region near the EP transposon insertion site within the first intron. We therefore conclude that forced PHF7 expression initiates an autoregulatory feedback loop which overcomes transcriptional repression of the testis-specific promoter.

**Fig.4.**
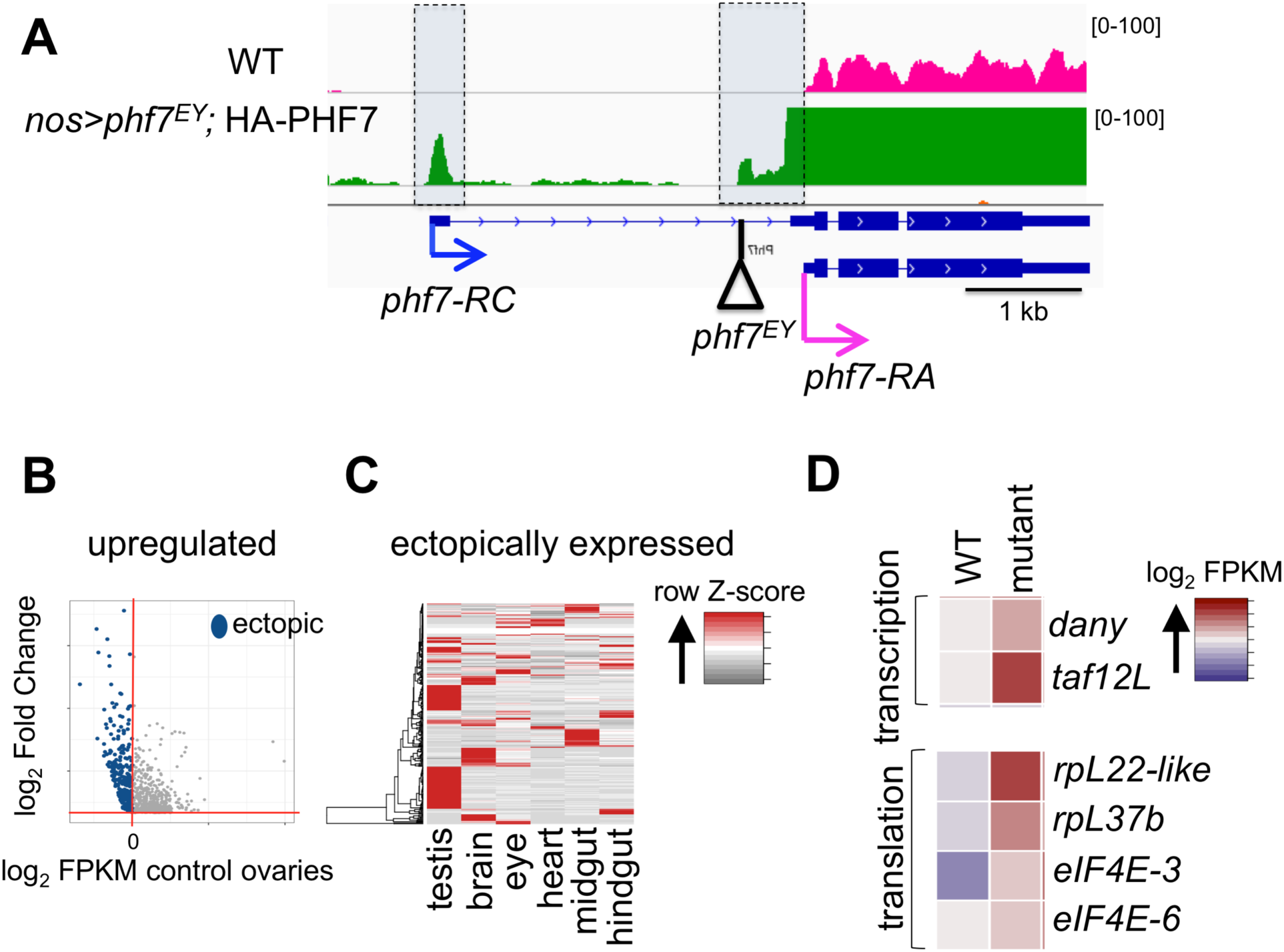
Ectopic *phf7* leads to female-to-male reprogramming of many genes including itself. (A) IGV genome browser view of the *phf7* locus. Wild-type RNA-seq reads are in pink and *nos*>*phf7*^*EY*^; *HA-phf7* RNA-seq reads are in green. The screen shot is reversed so that the start of transcription is on the left and all tracks are viewed at the same scale. Beneath is the RefSeq gene annotation of the two *phf7* transcripts and the location of the EP-element insertion in the endogenous locus of *phf7*^*EY*^. B) Scatter plots of significantly upregulated genes in *nos*>*phf7*^*EY*^; *HA-phf7* mutant ovaries. The log_2_ fold change in gene expression is plotted against the log_2_ of the FPKM values in wild-type ovaries. Blue points indicate ectopically expressed genes, genes which are not expressed in wild-type ovaries (log_2_ < 0). (C) Tissue expression clustering of the ectopically expressed genes in *nos*>*phf7*^*EY*^; *HA-phf7* mutant ovaries, displayed as a Z-score heatmap. Each column is an adult tissue. Each row is an ectopically expressed gene. (D) Expression values of select testis-specific genes in wild-type ovaries and *nos*>*phf7*^*EY*^; *HA-phf7* mutant ovaries displayed as a heat map of log2 FPKM values from RNA-seq analysis.

### Ectopic PHF7 provokes testis-specific gene expression programming

To gain a genome-wide view of the expression changes downstream of ectopic PHF7 expression we compared the transcriptomes of mutant ovaries with wild-type ovaries from newborn (0-24 h) females. This analysis identified 835 genes that were downregulated at least 2-fold (FDR <0.05; Table S1), and 799 genes that were upregulated at least 2-fold (FDR <0.05; Table S2) in *nos*>*phf7*^*EY*^; *HA-phf7* mutant ovaries. 286 of the upregulated genes are not detectable in wild-type ovaries (FPKM <1; Fig. 4B). Hierarchical clustering of the ectopically expressed genes revealed a tissue-specific signature most similar to samples from adult testis (Fig. 4C). Indeed, 60% (173/286) of the ectopically expressed tumor genes are genes known to be highly expressed in normal testis. Interestingly, several of the aberrantly expressed testis genes encode components of the specialized transcription and translation machinery that operates in the male germline (White-Cooper and Caporilli, 2013). These include Taf12L, Dany, RpL37b, RpL22-like, eIF4E-3, and eIF4E-6 (Fig. 4D). The acquisition of these male germ cell features reveals that forcing PHF7 expression leads to a loss of sexual identity and adoption of a male fate.

Given that ectopic expression of PHF7 in ovarian germ cells can drive transcription of normally silenced male germ cell genes, we hypothesized that PHF7 controls expression of the same set of genes in male germ cells. In an analysis focused on *phf7* function in the male germline, only 45 genes were found to be differentially expressed in *phf7* loss of function testis (Yang et al., 2017). Only one of these genes (CG15599) is ectopically expressed in PHF7-expressing ovarian germ cells. An additional 16 genes are significantly affected in both PHF7-expressing ovarian germ cells and *phf7* loss of function male germ cells (Table S3). Thus, contrary to our expectations, we found that the majority of genes under PHF7 control are different in male and female germ cells. These observations imply that tissue-specific factors define the genes and pathways under PHF7 control and raises the possibility that PHF7’s role in female germ cells may be different from its role in male germ cells.

In summary, our studies highlight the importance of preventing expression of lineage-inappropriate genes for maintaining tissue homeostasis. We demonstrate that once expressed in female germ cells, PHF7 can increase its own expression via a positive autoregulatory feedback loop. Our data further suggest that high levels of ectopic PHF7 are prerequisite to activate a male germ cell transcriptional program that drives tumor initiation and progression in female germ cells. PHF7 is a presumed chromatin reader as it preferentially binds to H3K4me2 *in vitro*, a mark generally enriched at transcription start sites (Yang et al., 2012). However, the mechanism by which PHF7 controls gene expression *in vivo* has remained elusive due to the lack of experimentally confirmed target genes. We speculate that, in female germ cells, PHF7 redirects chromatin remodeling complexes to inappropriately activate male germ cells genes. Future studies will focus on identifying the genes directly controlled by PHF7 to reveal the mechanism by which PHF7 fuels the oncogenic gene expression program.

## MATERIALS AND METHODS

### Drosophila stocks and culture conditions

The wild-type reference strain was *y*^*1*^, *w*^*1*^ (BDSC #1495). The following stocks were used to ectopically express *phf7* in female germ cells: P{GAL4::VP16-nos.UTR} (BDSC #4937), P{EPgy2}Phf7^EY03023^ (BDSC #15894), P{UASz-Phf7} (this study) and P{UASz-Phf7^ΔPHD^} (this study). The following stocks were used to report on *phf7* gene activity: The HA-tagged *phf7* transgene, PBac{3XHA-PHF7}, generated in the Van Doren lab by tagging the *phf7* locus included in the 20 kb BAC construct CH322-177L19 and inserted it into the 65B2 PBac{y[+]-attP-3B}VK00033 site via *phi-C31* catalyzed integration (Yang et al., 2012), and the *phf7*^*ΔORF*^::*GFP* allele (this study).

Drosophila stocks were maintained at 25°C. Crosses to drive ectopic expression with the UASz transgenes were set up at 29°C and the adults were aged 3-5 days prior to gonad dissection. Crosses to generate ectopic expression with P{EPgy2}Phf7^EY03023^ were set up at 18°C and the adults were transferred to 29°C for 10 days prior to gonad dissection.

### Generation of transgenic lines

Constructs were generated in the UASz expression vector to maximize expression in the female germline (DeLuca and Spradling, 2018). The P{UASz-Phf7} transgene was constructed by cloning the *phf7*cDNA (LD43541) into the mini-white containing *pUASz1.1* transformation vector (DGRC #1433). To construct the P{UASz-Phf7^ΔPHD^} transgene, an NEB Q5 mutagenesis kit was used to generate a deletion in the coding region using the primers: F-5’-TGCCATCAGCATGTGCTG-3’ and R-5’-AAGCAAACGGCAGCGGTT-3’. The transgenic constructs were sent for *phi-C31* catalyzed integration into the 65B2 PBac{y[+]-attP-3B}VK00033 site (Rainbow Transgenic Flies, Inc).

### Generation of the *phf7*^*ΔORF*^*-GFP* allele

The *phf7*^*ΔORF*^::*GFP* allele was generated using CRISPR to replace the *phf7* open reading frame with GFP. To generate the *phf7*^*ΔORF*^ deletion allele, the following guide RNAs were synthesized and ligated into the pU6-BbsI-chiRNA vector (Addgene #45946):

gRNA1: F-5’- CTTCGGTCACCGGAAACGCATCCA-3’ and

R-5’- AAACTGGATGCGTTTCCGGTGACC-3’.

gRNA2: F-5’- CTTCGAATCCTTGCGGCTGGCCATG-3’ and

R-5’- AAACCATGGCCAGCCGCAAGGATTC-3’.

1 kb homology arms were generated through PCR and cloned into the pHD-dsRed-attP (Addgene #51019). Guide RNAs and the donor vector were co-injected into *vas-Cas9* embryos (BDSC #51324; Rainbow Transgenic Flies, Inc). To insert GFP, the *gfp* coding sequence was cloned into the attB containing RIV-white^+^ transformation vector (DGRC #1330) and sent for *phi-C31* catalyzed integration into the attP site present in *phf7*^*ΔORF*^ (Rainbow Transgenic Flies, Inc).

### Immunofluorescence and image analysis

Ovaries and testis were fixed and stained according to standard procedures with the following primary antibodies: rat anti-HA (1:500, Roche cat# 11867423001, RRID: AB_390919), rat anti-Vasa (1:100, Developmental Studies Hybridoma Bank, RRID: AB_760351) and rabbit anti-GFP (1:2500, Thermo Fisher, cat# A-11122, RRID: AB_221569). Staining was detected with the following conjugated antibodies: Fluorescein (FITC) anti-rat (1:200, Jackson ImmnoResearch Labs, cat#A-21434, RRID: AB_2535855), FITC anti-rabbit (1:200, Jackson ImmnoResearch Labs, cat#111-095-003, RRID: AB_2337972), Alexa Fluor 555 anti-rat (1:200, Thermo Fisher, cat# A-21434, RRID: AB_2535855) or Alexa Fluor 555 anti-rabbit (1:200, Thermo Fisher, cat# A-21428, RRID: AB_2535849). TO-PRO-3 Iodide monomeric cyanine nucleic acid stain (Thermo Fisher, cat# T3605) was used to stain DNA.

Images were taken on a Leica SP8 confocal with 1024×1024 pixel dimensions, a scan speed of 600 Hz, and a frame average of 3. Sequential scanning was done for each channel and three Z-stacks were combined for each image. Processed images were compiled with Gnu Image Manipulation Program (GIMP) and Microsoft PowerPoint.

### qRT-PCR and data analysis

RNA was extracted from dissected gonads using TRIzol (Thermo Fisher, cat# 15596026) and DNase RQ1 (Promega, cat# M6101). Quantity and quality were measured using a NanoDrop spectrophotometer. cDNA was generated by reverse transcription using a SuperScript First-Strand Synthesis System for RT-PCR Kit (Thermo Fisher, cat# 11904018) using random hexamers. qPCR was performed using Power SYBR Green PCR Master Mix (Thermo Fisher, cat# 4367659) with an Applied Biosystems 7300 Real Time PCR system. PCR steps were as follows: 95°C for 10 minutes followed by 40 cycles of 95°C for 15 seconds and 60°C for 1 minute. Melt curves were generated with the following parameters: 95°C for 15 seconds, 60°C for 1 minutes, 95°C for 15 seconds, and 60°C for 15 seconds. Measurements were taken in biological triplicate with two technical replicates each. Relative transcript levels were calculated using the 2-ΔΔCt method (Livak and Schmittgen, 2001).

To measure RNA levels in *phf7*^*ΔORF*^::*GFP* gonads, the primer sequences were: for *phf7-RC*, F-5’-AGTTCGGGAATTCAACGCTT-3’ and R-5’-GAGATAGCCCTGCAGCCA-3’; for *gfp*, F-5’-ACGTAAACGGCCACAAGTTC and R-5’-AAGTCGTGCTGCTTCATGTG-3’.

To measure RNA levels in wild-type and *phf*^*EY*^ gonads, the primer sequences were for *phf7-RC, phf7-RC*, F-5’-AGTTCGGGAATTCAACGCTT-3’ and R-5’-GAGATAGCCCTGCAGCCA-3’; for total *phf7* F-5’-GAGCTGATCTTCGGCACTGT-3’ and R-5’-GCTTCGATGTCCTCCTTGAG-3’.

To measure RNA levels from the PBac{3XHA-PHF7} transgene, the primers were for *HA-phf7-RC*, F-5’-CTGCAGGGCTATCTCCGATA-3’ and R-5’-TAGCCCGCATAGTCAGGAAC -3’; for total *HA-phf7*, F-5’-CGATGTTCCTGACTATGCGG-3’ and R-5’-ACAGTGCCGAAGATCAGCT-3’

### RNA-seq and data analysis

Total RNA was extracted from dissected ovaries using TRIzol (Thermo Fisher, cat# 15596026). RNA quality was assessed with Qubit and Agilent Bioanalyzer. Libraries were generated using the Illumina TruSeq Stranded Total RNA kit (cat# 20020599). Sequencing was completed on 2 biological replicates of each genotype with the Illumina HiSeq 2500 v2 with 100bp paired end reads. Sequencing reads were aligned to the *Drosophila* genome (UCSC dm6) using TopHat (2.1.0) (Trapnell et al., 2009). Differential analysis was completed using CuffDiff (2.2.1) (Trapnell et al., 2012). Genes were considering differentially expressed if they exhibited a two-fold or higher change relative to wild-type with a False Discovery Rate (FDR) <0.05. The wild-type ovary mRNA-seq data sets are available from the National Center for Biotechnology Information’s GEO database under accession number GSE109850. and the mutant ovary mRNA-seq data sets are available under accession number GSE150213.

The screen shot of the expression data is from Integrated Genome Viewer (IGV). To account for the differences in sequencing depth when creating IGV screenshots, the processed RNAseq alignment files were scaled to the number of reads in the wild type file. This was done with Deeptools bigwigCompare using the scale Factors parameter with a bin size of 5.

Scatter plots were generated using ggplot function in R. Genes that were expressed in mutant (FPKM ≥ 1) but not expressed in wild type ovaries (FPKM<1) were called ectopic.

Tissue expression clustering of the ectopically expressed genes was performed to identify tissue-specific signatures. Expression values normalized to the whole fly were extracted from FlyAtlas (versions 1 and 2) (Leader et al., 2018; Robinson et al., 2013). The heatmap to compare the tissue expression profile of these genes per tissue was generated in R with heatmap.2 (gplots). Genes were clustered and normalized per row.

## Acknowledgments

We would like to thank Dr. Mark Van Doren, the Bloomington Drosophila Stock Center and the Iowa Developmental Studies Hybridoma Bank for fly stocks and antibodies. We would like to thank the Genomics Core at CWRU for performing the RNA-seq and Jane Heatwole for fly food. Imaging was performed using equipment purchased through NIH S10OD016164. This work was supported by the National Institutes of Health, R01GM129478 to H.K.S. and T32GM008056 to A.E.S.

## Data Availability

All raw and processed sequencing data generated during the course of this analysis can be found in GEO under accession number GSE150213.

## Figure Legends

**Fig. S1.**
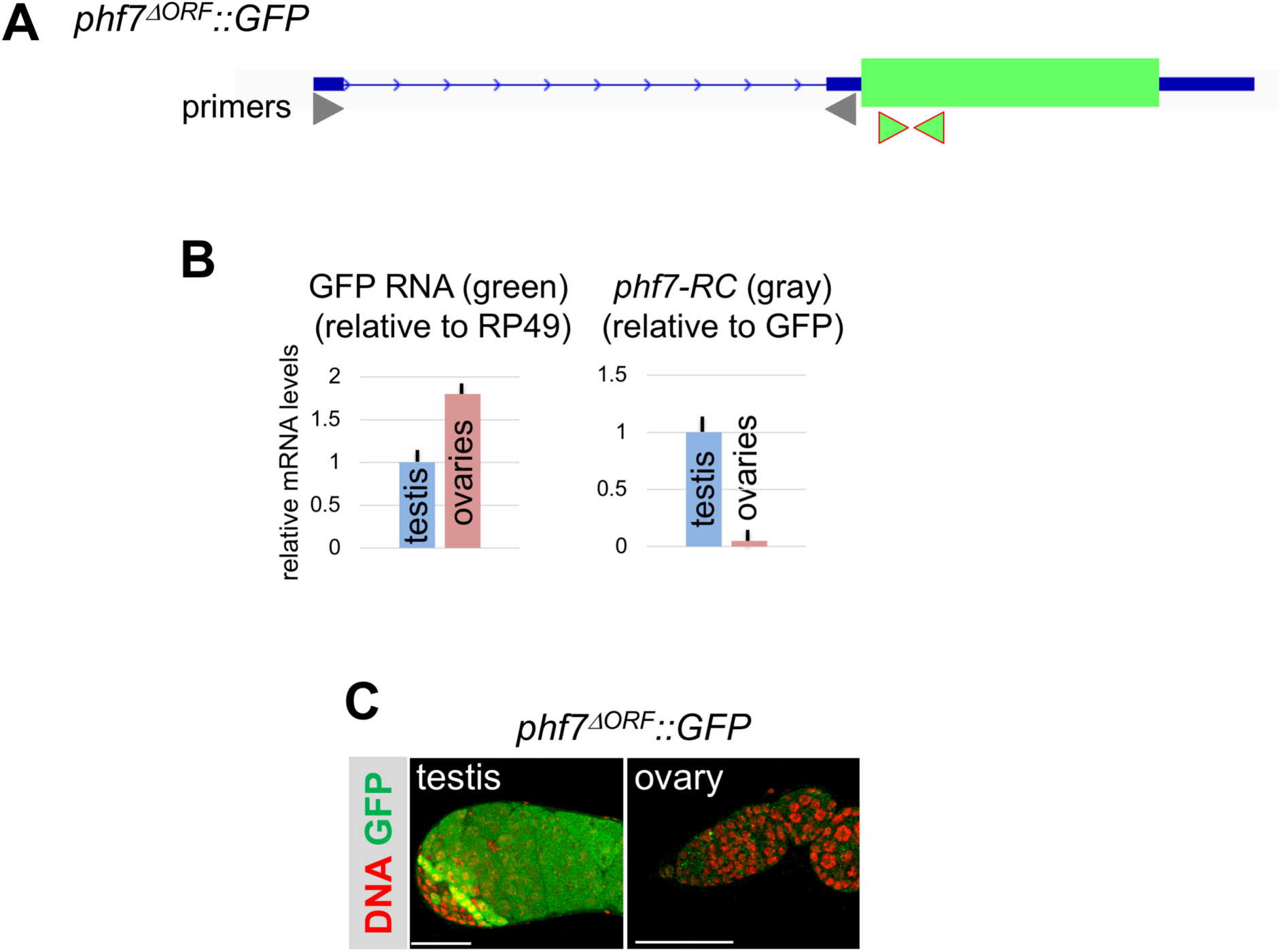
The *phf7*^*ΔORF*^::*GFP* allele is sex-specifically regulated. (A) Schematic of the *phf7*^*ΔORF*^::*GFP* allele in which the open reading frame was replaced by GFP (green box). Primers for RT-qPCR are indicated by arrowheads: Gray for *phf7-RC*, and Green for GFP. (B) RT-qPCR measurements of *phf7-RC* transcript in *phf7*^*ΔORF*^::*GFP* ovaries and testis. Error Bars indicate standard deviation of three biological replicates. (C) Confocal images of testis from hemizygous *phf7*^*ΔORF*^::*GFP* males and ovaries from homozygous *phf7*^*ΔORF*^::*GFP* females stained for GFP protein (green) and DNA (red). Scale bar, 50 μm.

